# Intra-molecular Electro-potential Circuit & ElectroNegatrode: Hypothesis, Algorithm & Implementation for universal indicative rule towards activity of biomolecules

**DOI:** 10.1101/2020.03.05.979807

**Authors:** Om Prakash

## Abstract

No universal indicative rule has been defined till now for activity & selectivity of biomolecules. This lacks very early perception about any molecule (specifically small molecule) in the biological system, beyond doing any experiment. The present study is all about hypothesizing, algorithm development and implementation of theoretical processing of 3D structure of biomolecules in the search of ground for developing perception about biomolecules individually or in combination of other biomolecules. The developed algorithm has been evaluated with small molecules and was found to be implementable as universal indicative rule towards activity & selectivity of biomolecules.

## INTRODUCTION

In present study, efficacy of intra-molecular information has been exploited for observation of molecules in biological systems. Recently, kinetics of intra-molecular diffusion of other molecules has been studied for understanding the effect of transfer of electron on catalysis (Del Barrio et al., 2018). Concept of open circuit potential has been observed in concern of organic substances for elevation of voltage (Sil et al., 2018)(Cui et al., 2019). Importance of location, size and orientation were discussed in concern of intermolecular junctions (Yao et al., 2007). Relations of effect of mass and intermolecular transport of charges were discussed (Yang et al., 2017). Existence and importance of auto regulatory circuits has been already postulated in various signaling networks as p53 & NF-kB etc. Effects of such circuits were also kept in consideration in reference of various phenotypic observations (Salminen and Kaarniranta, 2011). Formation of bond and cyclic charging & discharging has also been notified under the influence of change in potentials in the circuits (Motoyama et al., 2017). Concept of circuit has also been defined in relation of auto regulation of inflammatory responses, and its effect on molecular lytic process due to transfer of charges (Mancini and Di Battista, 2011). Concept of intramolecular charge transfer has also been adopted for development of small molecular devices (Yao et al., 2017). Relation between structure and its properties has been rationalized by using intermolecular transfer of charge and potential (Hasanein et al., 2016). Generation of energy of specific capacity was also discussed in relation of density of potential and circuits (Ren et al., 2016). Effect of specific amino acid residue and utility of intramolecular bonds were related for responding towards activity on other organisms (Himmerkus et al., 2010). In another study binding capacity was also related with electrochemical properties (Gong and Li, 2011). Beyond all these, a theoretical characterization of intramolecular potential has also been attempted for designing of small molecule material using organic molecules (Duan et al., 2013). Long range charge distribution for signal transfer was exploited with intramolecular charge distribution in neighboring objects. This study was concluded also into the cellular automata (Lu and Lent, 2011).

Being inspired from the influential utility of intramolecular information, in present study intramolecular EP-circuit & ElectroNegatrode has been hypothesized. Since no theoretical basis for universal indicative rule towards activity of biomolecules has been defined, therefore attempts has been made to use the intramolecular information for describing universal indicative rule towards activity of biomolecules.

## MATERIAL & METHOD

Every atom has its polarity. It is hypothesized that every molecule can be represented as meta-polarity of atoms. This meta-polarity can be constructed by sequential association and dissociation of Pauling electro-negativity of atoms. Pauling electronegativity has been used in various studies as: reactivity of reactions (Yu et al., 2018), in relating quantum mechanics with classical molecular dynamics (Santos et al., 2019), and understanding the effect on bond strength due to intramolecular electron delocalization (Lin et al., 2018) etc. Algorithm has been defined for this process (Figure 1). Since algorithm proceeds through a sequential process and ultimately electronegative poles (here called Electronegatrode) came in existence, therefore it has been defined as intramolecular electro-potential (EP) circuit. Extent of closeness of terminals EP-circuit has been defined with distance between electronegatrodes. Extent of closeness of EP-circuit has been evaluated in relation of activity & selectivity of biomolecules. This evaluation has been performed with small molecules, protein molecules and complex molecules. After evaluation, it was found that EP-circuit based observations can be directly used as theoretical basis for universal indicative rule towards activity & selectivity of biomolecules. Different derivatives of hypothesis of EP-circuit can be defined for various purposes related with molecules.

**Figure 1.**
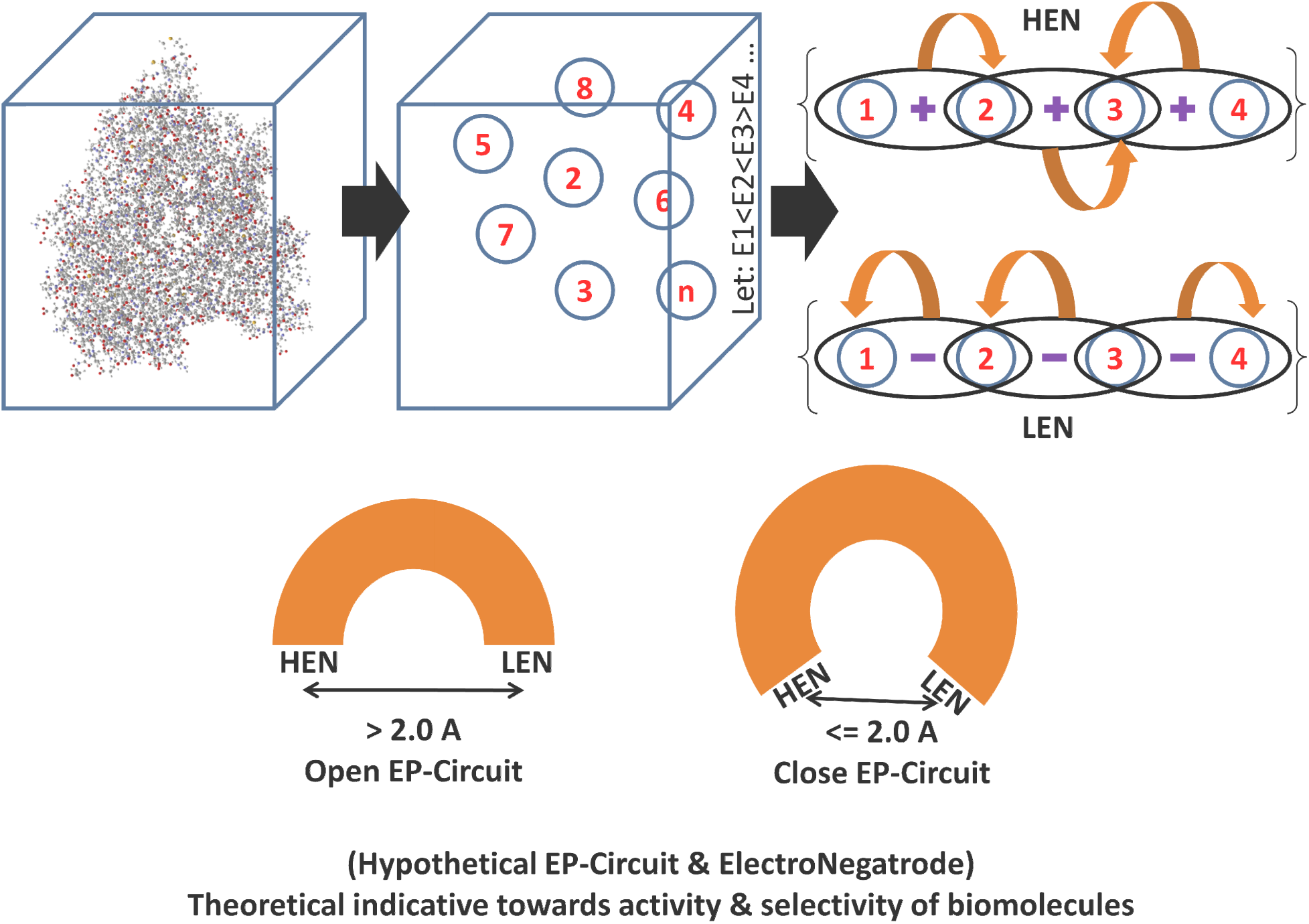
Pictorial representation of hypothesis of intramolecular EP-circuit

### Molecule observation criteria

After defining the intramolecular EP-circuit in biomolecules, it should be observed on the basis of open & closed circuit. This has been defined by distance (in Å) between terminals of circuit. Closed EP-circuit has been considered with <= 2Å distance between terminals; while open circuit will have this distance as > 2Å.

### Algorithm evaluation on biological system

Existence of importance of distance between terminals of EP-circuit, has been evaluated as classification capability. For evaluation, the developed algorithm has been implemented with GABA receptor binders as active (K_i_ < 10000 nM) & inactive (K_i_ >= 10000 nM) classes.

## RESULTS & DISCUSSION

### Pseudo code for algorithm in the search intrmolecular EP-circuit

PDB file including biomolecule(s) is starting input to algorithm. PDB file contained location of atoms of molecule in three dimensions. 3D finite dimensional space (*I*^*3*^) was presented in matrix *C* of a × 3, where three columns presents X, Y, & Z coordinates; and a represented number of atoms in the molecule. *I*^*3*^ was further exploited into 2D matrix R of order a × 2, where R include two columns of Pauling Electronegativity value and Atomic weight for each atom.

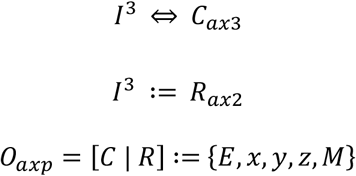

Let origin at (0, 0, 0) coordinates of *I*^*3*^ then:

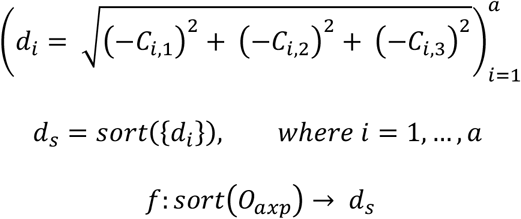

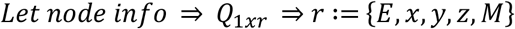

### For High Electronegativity Node

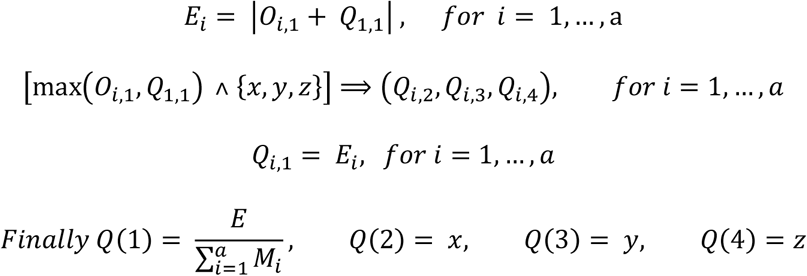

### For Low Electronegativity Node

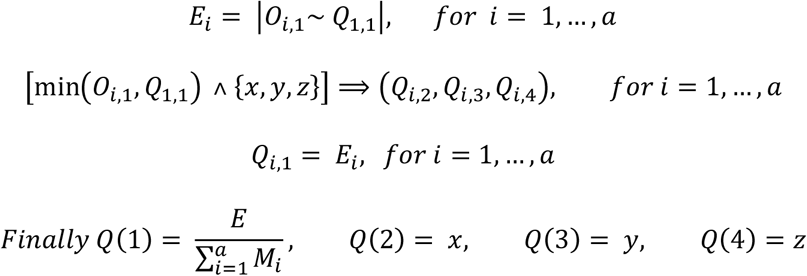

### Molecule observation criteria

Intramolecular EP-circuit bears two terminals (here called ElectroNegatrode), one with High Electronegativity value per unit mass (H_EN_) and other with Low Electronegativity value per unit mass (L_EN_). Closeness & Openness of EP-circuit was defined on the basis of distance between two terminals of EP-circuit.

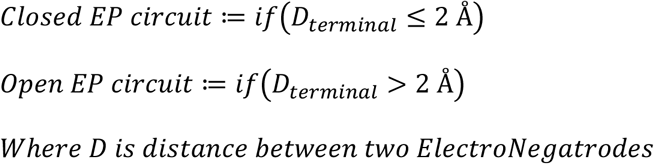

### Evaluation of hypothesis on the basis of classification of molecules

By considering the criteria for open & closes circuits and effectiveness of hypothesis for defining a perception about any molecule in biological system; threshold of 2Å was used for evaluation of 68 small molecules known experimentally against GABA receptors. On the basis of K_i_ value of small molecules against GABA receptors, two classes were defined as Active (K_i_ < 10000 nM) and Inactive (K_i_ >= 10000 nM). First 24 compounds (G1-24) belonged to active class; and remaining 44 were considered into inactive class. Classification capacity of present hypothesis was evaluated on data.

|  | Correct Prediction | Wrong Prediction |
| --- | --- | --- |
| Active (24) | 15 (TP) | 09 (FN) |
| Inactive (44) | 33 (TN) | 11 (FP) |

TP: True Positive; TN: True Negative; FP: False Positive; FN: False Negative

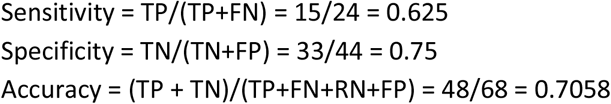

Classification statistics favored the efficiency of hypothesis of intramolecular EP-circuit & ElectroNegatrode for developing early perception about any small molecule.

## CONCLUSION

After development and evaluation of algorithm for Intra-molecular Electro-potential Circuit & ElectroNegatrode, hypothesis of EP-circuit can be further exploited for development of universal indicative rules towards activity of biomolecules, including extent of closed level of EP-circuit.

## ACKNOWLEDGEMENT

Author express gratitude to *The Institute of Mathematical Sciences*, Chennai-600113, India for providing research facilities as well as DAE Post-Doctoral Fellowship (PDF 214). Author is also thankful to Dr. Amit Singh (PDF Mathematics, IMSc Chennai) for mathematical proof editing.

